# Inter-Subject Correlation during New Music Listening: A Study of Electrophysiological and Behavioral Responses to Steve Reich’s *Piano Phase*

**DOI:** 10.1101/2021.04.27.441708

**Authors:** Tysen Dauer, Duc T. Nguyen, Nick Gang, Jacek P. Dmochowski, Jonathan Berger, Blair Kaneshiro

## Abstract

Musical minimalism utilizes the temporal manipulation of restricted collections of rhythmic, melodic, and/or harmonic materials. One example, Steve Reich’s *Piano Phase*, offers listeners readily audible formal structures containing unpredictable events at local levels. Pattern recurrences may generate strong expectations which are violated by small temporal and pitch deviations. A hyper-detailed listening strategy prompted by these minute deviations stands in contrast to the type of listening engagement typically cultivated around functional tonal Western music. Recent research has suggested that the inter-subject correlation (ISC) of electroencephalographic (EEG) responses to natural audio-visual stimuli objectively indexes a state of “engagement”, demonstrating the potential of this approach for analyzing music listening. But can ISCs capture engagement with minimal music, which features less obvious expectation formation and has historically received a wide range of reactions? To approach this question, we collected EEG and continuous behavioral (CB) data while 30 adults listened to an excerpt from Steve Reich’s *Piano Phase*, as well as three controlled manipulations and a popular-music remix of the work. Our analyses reveal that EEG and CB ISC are highest for the remix stimulus and lowest for our most repetitive manipulation. In addition, we found no statistical differences in overall EEG ISC between our most musically meaningful manipulations and Reich’s original piece. We also found that aesthetic evaluations corresponded well with overall EEG ISC. Finally we highlight co-occurrences between stimulus events and time-resolved EEG and CB ISC. We offer the CB paradigm as a useful analysis measure and note the value of minimalist compositions as a limit case for studying music listening using EEG ISC. We show that ISC is less effective at measuring engagement with this minimalist stimulus than with popular music genres and argue that this may be due to a difference between the type of engagement measured by ISC and the particular engagement patterns associated with minimalism.

## 1 Introduction

The genre of musical minimalism is famously (or, perhaps infamously depending on the listener) characterized by highly recurrent, starkly restricted pitch and rhythmic collections. From the early days of scholarship on minimal, or “repetitive music” as it was often called, commentators described the music’s timbral and rhythmic staticity and its limited pitch patterns (Mertens, 1983, p. 12). While many advocates reported what we might call blissing out to this “meditative music” (to use yet another early term for this repertoire), some composers went on record to state their intention that the music should be listened to carefully (Henahan, 1970; Strongin, 1969). For example, the composer Steve Reich wrote in 1968 that he wanted to write works with musical processes that listeners could perceive: works where the process unfolded very gradually in order to “facilitate closely detailed listening” (Reich, 2009, p. 34). Reich’s *Piano Phase* (1967) shows how this type of granular listening might unfold. The piece, written for two pianos or marimbas, alternates between two distinct and highly repetitive states resulting from a single process. During in-phase sections, the two performers play a short musical unit in rhythmic unison, though varying in pitch alignment (Figure 1). In between these in-phase sections, one performer gradually accelerates, resulting in unpredictable note onsets (i.e., phasing sections). Over time these phasing sections lead to a new pitch alignment in the subsequent in-phase section.^1^ The driving phasing process offers the listener an outline of how the piece unfolds at a macro-level while leaving many details unpredictable—from rhythms during the phasing sections to accent patterns during in-phase sections. For a listener interested in detailed minutia and slight variation, the work may fascinate; in other moods or with other priorities, the piece can bore, confuse, and even anger (Rockwell, 1973). How might we measure listeners’ engagement with such repertoire, given its reduced musical parameters and varied and polarized reception (Dauer, 2020)?

**Figure 1:**
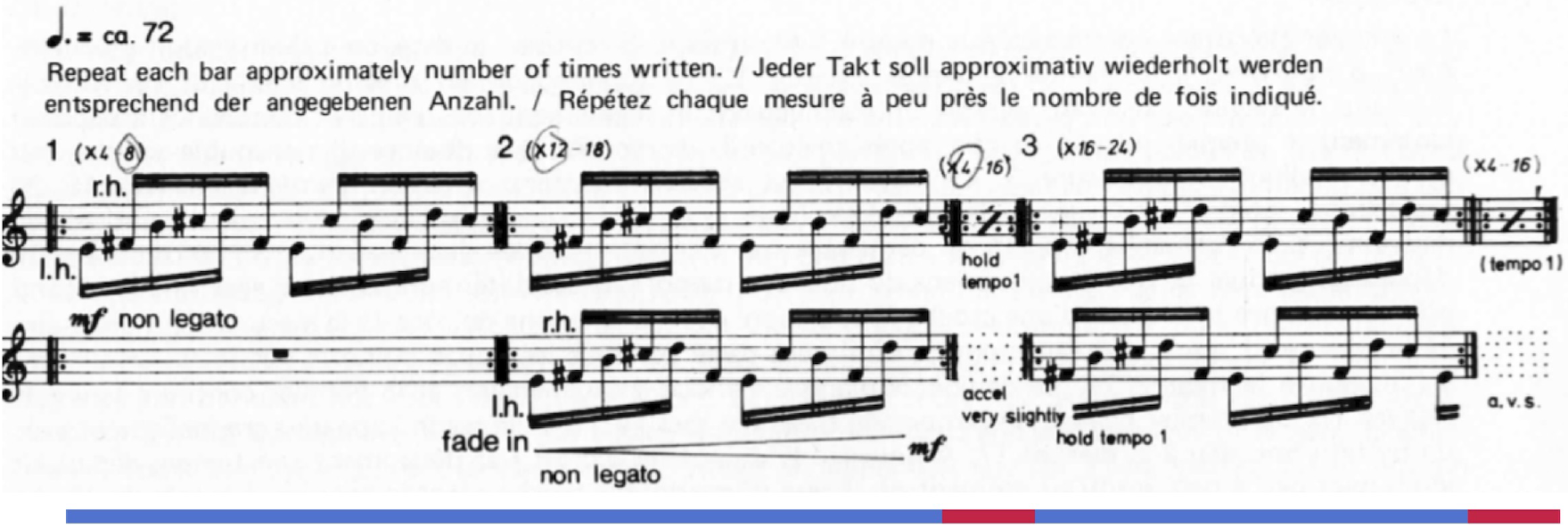
The opening modules from Steve Reich’s *Piano Phase*. Lines under the staff indicate sections: blue lines are in-phase sections and red lines are phasing sections.

Recent research using the high temporal resolution of electroencephalography (EEG) has suggested that the correlation of neural responses among participants (inter-subject correlation, or ISC) in response to natural audio-visual stimuli objectively indexes a state of “engagement” (Dmochowski et al., 2012). Ensuing studies have extended this work to musical stimuli and demonstrated how ISC may be a powerful tool for analyzing listening (B. Kaneshiro et al., 2020; Madsen et al., 2019; B. Kaneshiro et al., 2021). B. Kaneshiro et al. (2020) presented popular, Hindi-language songs from “Bollywood” films to participants and reported higher behavioral ratings and ISCs for their original versions when compared with phase-scrambled manipulations. Madsen et al. (2019) drew on instrumental compositions (nineteen Western classical musical works in a variety of styles, and one Chinese folk song) to establish that ISCs decrease over repeated exposures to familiar music (though ISCs were sustained for participants with musical training). Most recently, B. Kaneshiro et al. (2021) investigated participants’ time-resolved ISCs in response to the first movement of Edward Elgar’s Cello Concerto in E minor, Op. 85. In contrast to the stimuli used in these previous studies, and true to minimalism’s stereotypical characteristics, Reich’s *Piano Phase* features a high level of repetition, unchanging timbre, and narrow pitch content.^2^

Our primary research question was to uncover whether participants shared engagement patterns (as measured by ISC) while listening to *Piano Phase*. In particular, we hypothesized that phasing sections would be more engaging (i.e., elicit more correlated responses) than in-phase sections, due to phasing sections’ rhythmic variety and unpredictability coupled with a wider variety of pitch interactions. If listeners deployed the hyper-detailed listening strategy described above, phasing sections would offer rich content with which to engage. On the other hand, detailed listening during phasing sections could lead to divergent patterns of engagement as listeners lock on to different dimensions or aspects of the music during these more eventful sections. Since ISC depends on time-locked similarities in neural data, these divergent but equally engaged listening styles may not result in significant correlations. To test this possibility, we introduced manipulations of *Piano Phase*. First, we created a version without phasing sections, anticipating that ISC would be lower for this manipulation if such phasing sections were being picked up in the original version. Second, listeners have also historically reported an arguably more mood-driven type of engagement with this type of music, which, in contrast with detailed listening, allows for a more internal floating away of attention, still connected to the stimulus but unlikely to be correlated between participants (Lloyd, 1966). Therefore, we also included a manipulation of *Piano Phase* with frequent changes in the content (resulting from reshuffling five-second segments of the original excerpt). If ISC indexes this style of engagement in *Piano Phase*, we predicted less of the listening style for this manipulation. To examine the possibility of listeners being bored by the original work, we also introduced a third control stimulus with extreme repetition, which we expected to elicit no meaningful engagement. Finally, we included a commercial remix of Reich’s original work in a popular style, which we conjectured would engage listeners and elicit EEG ISC comparably to previous experiments (B. Kaneshiro et al., 2020; B. B. Kaneshiro, 2016).

In line with recent work, we computed ISCs over entire excerpts and in shorter, overlapping time windows, giving us a sense of overall engagement as well as moment-to-moment patterns shared between audience members (Dmochowski et al., 2012; B. Kaneshiro et al., 2021). To provide complementary measures of what ISC is reliably indexing, participants rated the stimuli and additionally completed a second experimental block where they continuously reported their level of engagement with the stimuli. This allowed us to compare relationships for both overall and time-resolved neural and behavioral measures.

Other researchers have used minimalist compositions as experimental stimuli, similarly taking advantage of the works’ unusual musical properties. Musicologist Keith Potter and computer science colleagues used two early works by Philip Glass to compare information dynamics and musical structure (Potter et al., 2007). Psychologist Michael Schutz worked with percussionist Russell Hartenberger to examine desynchronization among performers of Reich’s *Drumming* (Hartenberger, 2016),^3^ and Daniel Cameron and colleagues have studied experiences of groove and neural entrainment using Reich’s *Clapping Music* (D. J. Cameron et al., 2019; D. Cameron et al., 2017). Dauer et al. (2020) examined preattentive cortical responses to various types of formal repetition using synthesized melodies based on early minimalist compositional techniques. The current study takes minimalism as an edge case in the applicability of neural correlation, uniting the repertoire’s extreme musical techniques (and unique reception history) with multivariate techniques for analyzing brain data.

## 2 Methods

### 2.1 Stimuli

All five stimuli in the experiment are related to Steve Reich’s *Piano Phase*, a much-anthologized example of American minimalism for two pianos or marimbas (Figure 1). In the experiment we used pianists Nurit Tilles and Edmund Neimann’s 1987 recording on the album *Reich “Early Works”* released by Double Edge (Reich, 1987). We used the first five minutes and five seconds (5:05) of the track’s 20:26 duration. We refer to this excerpt of *Piano Phase* used in the experiment as the *Original* condition (Figure 2A).^4^

**Figure 2:**
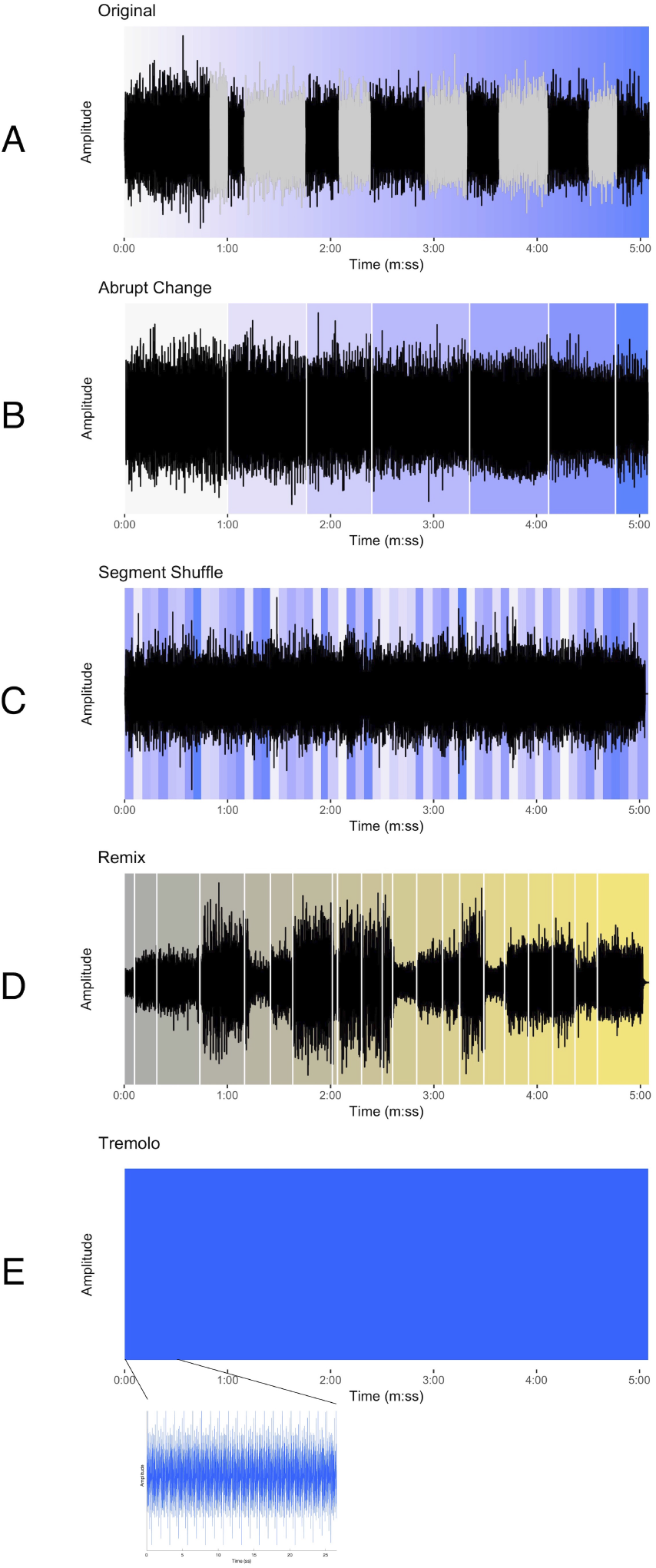
The waveforms for each of the stimuli in the experiment. (A) Original, with phasing sections colored gray and the progression of events represented by the gradual change of color from white to blue. (B) Abrupt Change, white lines denoting sudden shift from one in-phase section to the next and background color showing approximate location of in-phase material in the Original condition. (C) Segment Shuffle, random re-ordering of five-second units shown using original color in Original. (D) Remix (Winn’s *Piano Phase (D*Note’s Phased & Konfused Mix)*), gradual progression of events represented with color change from gray to yellow and key musical events beginning with white lines. (E) Tremolo, appearing as an unchanging block when zoomed out, but in the lower plot, zoomed in to show the reiterated pitch material.

*Piano Phase* is useful for exploring the limits of ISC in measuring musical engagement because it offers contrasting sections (phasing and in-phase) with slightly varying musical content for comparison while holding many other musical parameters constant: timbre, dynamics (largely), instrumentation, pitch content, and absence of lyrics or vocal content. These features make it uncommonly amenable to the creation of the stimulus manipulations used in this study.

Using MATLAB software, we created three stimulus conditions of equal duration, each based on the content of the excerpt used in the Original condition. First, in the *Abrupt Change* condition, (Figure 2B) all phasing sections from the Original excerpt were replaced with exact repetitions of the preceding in-phase material. The stimulus thus presents repetitions of an in-phase motif through the section where the phasing would have occurred, and then shifts abruptly to the next in-phase section as closely as possible to its occurrence in the original recording. For example, the stimulus begins with the in-phase section where Pianist 1 and Pianist 2 align the first notes of the twelve-note pattern. This continues without phasing until suddenly the next in-phase section emerges, where Pianist 2 aligns the second note of the pattern with the first note of the pattern played by Pianist 1. Thus, the Abrupt Change condition is, in essence, form without function: where regular markers of formal sections (i.e., points of arrival at the alignments of in-phase sections) are situated without the functional transitions (i.e., the phasing sections).

As a contrast to the sudden changes embodied by the Abrupt Change condition, we created the *Segment Shuffle* condition (Figure 2C). Here we divided the Original audio into five-second segments and randomly reordered them (i.e., “shuffled” them). In order to avoid abrupt disjunct shifts, the transitions between segments were smoothed by applying a linear crossfade. The five-second segments included both phasing and in-phase material, meaning that upcoming content was unpredictable for listeners. In contrast with the Abrupt Change condition, Segment Shuffle featured function without form: constant, potentially surprising changes with no overarching formal scheme.

Finally, we synthesized a stimulus with neither form nor function, taking the repetition aspect of minimalist music to an extreme. Our *Tremolo* condition (Figure 2E) consisted solely of the aggregated pitch content of *Piano Phase* presented as a block chord, reiterated at Reich’s opening tempo marking and lasting the duration of the Original excerpt.

For comparison with the more popular genres of audio material used in previous ISC studies, we also included Matt Winn’s *Piano Phase (D*Note’s Phased & Konfused Mix)*, an homage to Reich’s piece released on the 1999 *Reich Remixed* album (Reich, 1999); we refer to this condition as *Remix* for short. Winn’s dance music group, D*Note, draws on sounds from electronica and jazz, and these influences show up in Remix alongside samples from Reich’s piece.^5^ The entire track was used in the experiment and its duration (5:05) informed the length of the other stimuli. Listening to Remix, we identified moments (musical events) that we predicted would engage listeners (for a full list, see Table S1). These events guided our interpretation of time-resolved EEG and continuous behavioral (CB) results.

All stimuli were presented to participants as mono .wav files; the second audio channel was embedded with intermittent square-wave pulses which were used as precise timing triggers (see § 2.3 and B. Kaneshiro et al. (2020)).

### 2.2 Participants

We were interested in listeners’ initial experiences of Reich’s piece and sought participants who were unlikely to have heard the composition before. Participants had to be 18–35 years old, have normal hearing, be right-handed, have no cognitive or decisional impairments, be fluent in English, and have had no individual musical instrument or vocal training, nor musical education after high school (or equivalent).

The participant sample (*N* = 30; 19 female, 11 male) had a mean age of 23.8 years (ranging from 18 to 35 years). Twelve participants reported some formal musical training ranging from 2 to 16 years (average of 4.5 years) including activities such as elementary school band and orchestra and piano lessons in middle school. Only two participants reported ongoing musical activities (amateur ukulele playing and occasional jam sessions). All participants reported listening to music regularly, from 0.2 to 8 hours a day (average of 2.4 hours per day).

### 2.3 Experimental paradigm and data acquisition

The Stanford University Institutional Review Board approved this research, and all participants gave written informed consent before completing the experiment. After discussing and signing the consent form, each participant completed questionnaires about demographic information and musical experience. Each participant then completed two blocks: one EEG (Block 1) and one behavioral (Block 2), both conducted in an acoustically and electrically shielded ETS-Lindgren booth. The participant completed a brief training session to acquaint them with the interface and task before the experimenter donned the EEG net. The participant was told to sit comfortably in front of the monitor and view a fixation image While EEG was recorded. Participants listened to each of the five stimuli once in random order. Participants did not perform any task during the presentation of the stimuli and were told to refrain from moving their body in response to the music: they were told not to tap their feet or hands, or bob their heads. After each stimulus in Block 1, the participant rated how pleasant, well ordered, musical, and interesting the preceding stimulus was on a scale of 1 (not at all) to 9 (very) via key press using a computer keyboard. Participants were permitted to move and take short breaks in between stimuli (during which time a “break” screen appeared). When ready, the participant initiated the next stimulus by pressing the space bar on the keyboard.

The EEG net was removed after Block 1, and the participant returned to the sound booth to complete Block 2. Here the participant heard the same five stimuli (in random order) and this time completed a continuous behavioral task while listening. Their task was to continuously report their level of engagement—which was defined as “being compelled, drawn in, connected to what is happening, and interested in what will happen next” (Schubert et al., 2013)—over the duration of each stimulus. To perform this task, the participant used a computer mouse to control a slider shown on the computer monitor. After each stimulus, the participant rated how engaging they found the preceding stimulus to be overall, using the same 1–9 key press scale used in Block 1. The ordering of blocks was not randomized (i.e., the EEG block always preceded the CB block) because we wanted to ensure that during recording of EEG data in Block 1, participants would not be biased with the definition of engagement and the continuous reporting task that came in Block 2.

The experiment was programmed in MATLAB using the Psychophysics Toolbox (Brainard, 1997). Stimuli were played through two Genelec 1030A speakers located 120 cm from the participant. Stimulus onsets were precisely timed by sending square-wave pulses to the EEG amplifier from a second audio channel (not heard by the participant). We used the Electrical Geodesics, Inc, (EGI) GES 300 platform (Tucker, 1993), a Net Amps 300 amplifier, and 128-channel electrode nets to acquire data with a 1 kHz sampling rate and vertex reference. Before beginning the EEG block, we verified that electrode impedances were below 60 kΩ (Ferree et al., 2001).

### 2.4 EEG preprocessing

Continuous EEG recordings were preprocessed offline in MATLAB after export using Net Station software. The data preprocessing procedure used here is described in detail in B. Kaneshiro et al. (2021). Briefly, data were preprocessed on a per-recording basis: Each recording was highpass (above 0.3 Hz), notch (between 59 and 61 Hz) and lowpass (below 50 Hz) zero-phase filtered before being downsampled from 1 kHz to 125 Hz. Epochs for each stimulus were 5 minutes (37501 time samples) in length and precisely timed from the audio pulses. Ocular and EKG artifacts were removed using ICA (Jung et al., 1998), data were converted to average reference, and data from bad electrodes or noisy transients were replaced with a spatial average of data from neighboring electrodes. After preprocessing, each trial of data was a 2D electrode-by-time matrix (125 × 37501). The matrices contained data from 125 electrodes as we excluded the four sensors over the face (electrodes 125–128) and reconstituted the reference sensor during preprocessing (B. B. Kaneshiro, 2016; Losorelli et al., 2017; B. Kaneshiro et al., 2020, 2021). During preprocessing, participant S08’s response to the Tremolo stimulus was flagged as containing excessive noise artifacts; therefore we excluded this trial from further analysis, but retained other trials from this participant.

After preprocessing, we aggregated trials into 3D electrode-by-time-by-participant data matrices for each stimulus. As a result, responses to Original, Abrupt Change, Segment Shuffle, and Remix stimuli were stored in 125 × 37501 × 30 matrices, while responses to Tremolo were stored in a 125 × 37501 × 29 matrix.

### 2.5 Data analysis

Figure 3 summarizes our analysis pipeline for the EEG and CB data. EEG was recorded from participants in Block 1, and participants provided CB reports of engagement in Block 2. Participants also rated the stimuli in both blocks. We computed ISC of both the EEG and CB measures, and also computed mean CB across participants. Finally, we analyzed the ratings to determine whether they differed significantly according to stimulus.

**Figure 3:**
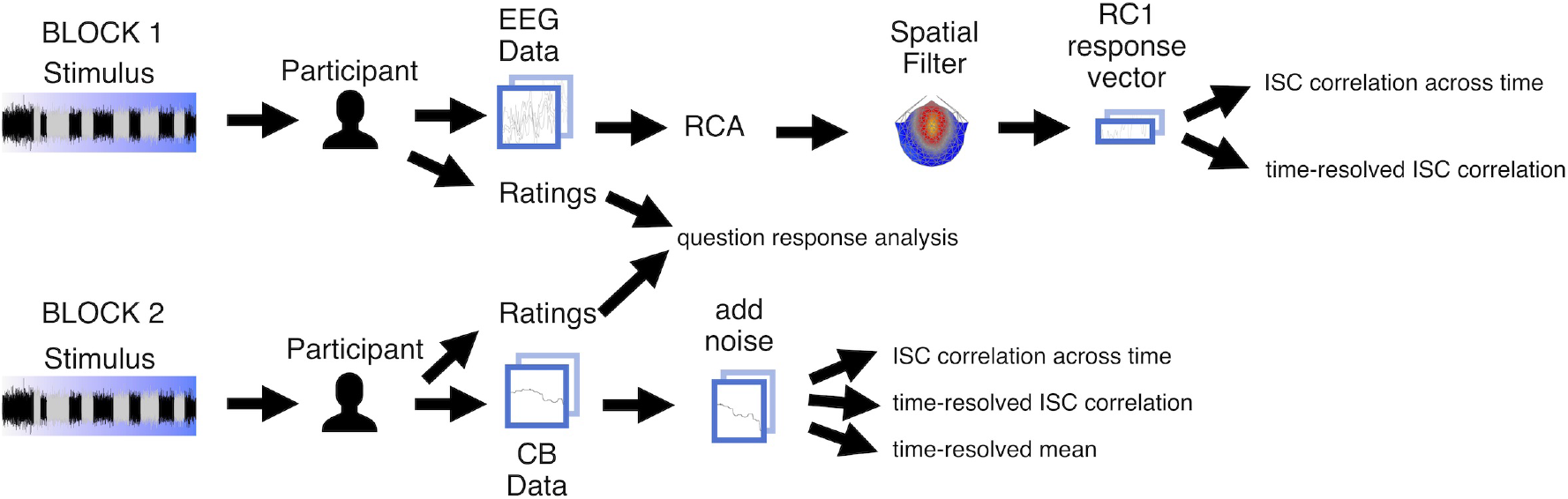
Analysis pipeline for experiment data. Participants heard each of the five stimuli twice, once in each block. During Block 1 we recorded EEG, and during Block 2 participants completed the continuous behavioral (CB) task. Participants answered questions about each stimulus after hearing it. For the EEG data we computed spatial components maximizing temporal correlation and projected electrode-by-time response matrices to component-by-time vectors. For vectorized EEG as well as CB vectors, we then computed inter-subject correlation (ISC) of the vectors on a per-stimulus basis, across time and in a time-resolved fashion. We additionally computed the time-resolved mean values between participants. We aggregated and analyzed ratings.

#### 2.5.1 Spatial Filtering of EEG Data

Previous EEG ISC studies have prepended a spatial filtering operation before computing correlations in order to maximize signal-to-noise ratio of the data while also reducing the dimensionality of each EEG trial from a space-by-time matrix to a time vector (Dmochowski et al., 2012). Therefore, we filtered the EEG data using Reliable Components Analysis (RCA) prior to computing ISC (Dmochowski et al., 2012, 2015). RCA maximizes across-trials covariance of EEG responses to a shared stimulus relative to within-trials covariance, and therefore maximizes correlated activity across trials (i.e., ISC). It is similar to PCA, but maximizes correlation across trials as opposed to variance explained in a single response matrix. Like PCA, RCA involves an eigenvalue decomposition of the data, and therefore returns multiple spatial filters as eigenvectors; and corresponding coefficients as eigenvalues (Dmochowski et al., 2012).

We used a publicly available MATLAB implementation (Dmochowski et al., 2015), computing RCA separately for each stimulus. Following B. Kaneshiro et al. (2020), we computed the top five reliable components. We observed a sharp drop in RC coefficients after the first, most-correlated component (RC1); given that past research has reported negligible ISC in subsequent RCs in this scenario (B. Kaneshiro et al., 2021), we proceeded with ISC analyses using RC1 data only, as was done by B. Kaneshiro et al. (2020). In presenting the component topographies on a scalp map, each weight vector was multiplied by ±1 such that frontal electrodes were associated with positive weightings; this was for visualization only, and polarity of the projected data does not impact computed correlations.

#### 2.5.2 Inter-Subject Correlation Analyses

We followed the procedure of B. Kaneshiro et al. (2021) to compute the EEG ISC of RC1 response vectors. First, we computed ISC across the entire duration of each stimulus. Following this, we computed ISC in a time-resolved fashion, over 5-second windows with a 1-second shift between windows. This gave us a total of 296 time-resolved ISC points across each stimulus with a temporal resolution of 1 second. ISC for each participant was computed in a one-against-all fashion (the correlation of each participant’s RC1 response vector with every other participant’s response vector for a given stimulus). We report the mean ISC across participants and additionally visualize single-participant correlations for all-time ISC and standard error of the mean for time-resolved ISC.

For the CB responses, we computed mean CB at each time sample, as well as CB ISC both across entire excerpts and in the short time windows described above. CB responses were already in vector form for each participant, so we did not perform any operation akin to EEG spatial filtering before computing means and ISC. At times, individual participants did not move the slider in a given five-second window, which would produce missing values when computing correlations. To address this issue, for the CB ISC analyses *only* we added a small amount of noise, uniformly distributed over the interval ±0.001, independently to each CB response vector prior to computing ISC. As with the EEG data, we report means and single-participant values for analyses across entire stimuli, and means with standard error of the mean for time-resolved measures.

#### 2.5.3 Statistical analyses

Significance of each EEG result was computed using permutation testing. As described in detail in previous studies (B. Kaneshiro et al., 2020, 2021), we conducted each EEG analysis 1,000 times; in each iteration, the phase spectrum of each EEG trial input to RCA had been randomized (Prichard & Theiler, 1994). The distribution of 1,000 outcomes for each analysis then served as the null distribution for assessing significance of the observed result. We performed a similar procedure to create null distributions for CB ISC, independently phase scrambling each CB response vector prior to computing ISC—also over 1,000 iterations.

Behavioral ratings, EEG ISC computed over entire stimuli, and CB ISC computed over entire stimuli were each analyzed using R (Ihaka & Gentleman, 1996; R Core Team, 2019) and lme4 (Bates et al., 2012). Here we performed a linear mixed-effects analysis of the relationship between response values and stimulus conditions, with fixed effect of condition (Original, Abrupt Change, Segment Shuffle, Remix, and Tremolo) and random effect of participant in each model. As in B. Kaneshiro et al. (2020), ordinal behavioral ratings were treated as approximately continuous (Norman, 2010). Following this we conducted two-tailed pairwise t-tests to assess differences between pairs of stimulus conditions.

Results for analyses involving multiple comparisons were corrected using False Discovery Rate (FDR, Benjamini & Yekutieli (2001)). For discrete results, we corrected for multiple comparisons on a per-stimulus basis (EEG ISC and CB ISC data: ten paired comparisons over five stimulus conditions; behavioral ratings: ten paired comparisons per stimulus; RC coefficients: five unpaired comparisons per stimulus). We performed no temporal cluster correction on the time-resolved ISC: as noted by B. Kaneshiro et al. (2021), temporal dependence was accounted for in the phase-scrambling procedure underlying the permutation testing, which preserves autocorrelation characteristics of the original response data (Prichard & Theiler, 1994; Lancaster et al., 2018).

## 3 Results

In order to examine engagement with an example of musical minimalism, we used intersubject correlation (ISC) to analyze EEG and continuous behavioral (CB) responses from 30 adult participants who heard an intact excerpt of Steve Reich’s *Piano Phase*, three manipulated control stimuli, and a professional remix of Reich’s piece. We analyzed EEG and CB ISC in two ways: an aggregate ISC value for each stimulus (full-time EEG ISC, full-time CB ISC) and time-resolved ISCs for both EEG and CB data. Each participant also gave ordinal ratings of each stimulus (behavioral ratings).

### 3.1 Remix Stimulus Garnered Highest Behavioral Ratings

After hearing each stimulus in Block 1, participants used a 1–9 scale to rate how pleasant, musical, well ordered, and interesting they found each excerpt. Later, in Block 2, they used the same scale to report their overall level of engagement with each stimulus. Based on a repeated-measures ANOVA, ratings for all five questions were found to differ significantly by condition (Figure 4): pleasant (χ^2^(4) = 126.03, *p* < 0.001), musical (χ^2^(4) = 139.78, *p* < 0.001), well ordered (χ^2^(4) = 37.996, *p* < 0.001), interesting (χ^2^(4) = 104.29, *p* < 0.001), and engaging (χ^2^(4) = 127.92, *p* < 0.001). Note that here we present the statistics in a question-wise fashion in order to emphasize differences between stimuli, in Figure 4 we grouped question responses by stimulus to emphasize patterns within stimuli.

**Figure 4:**
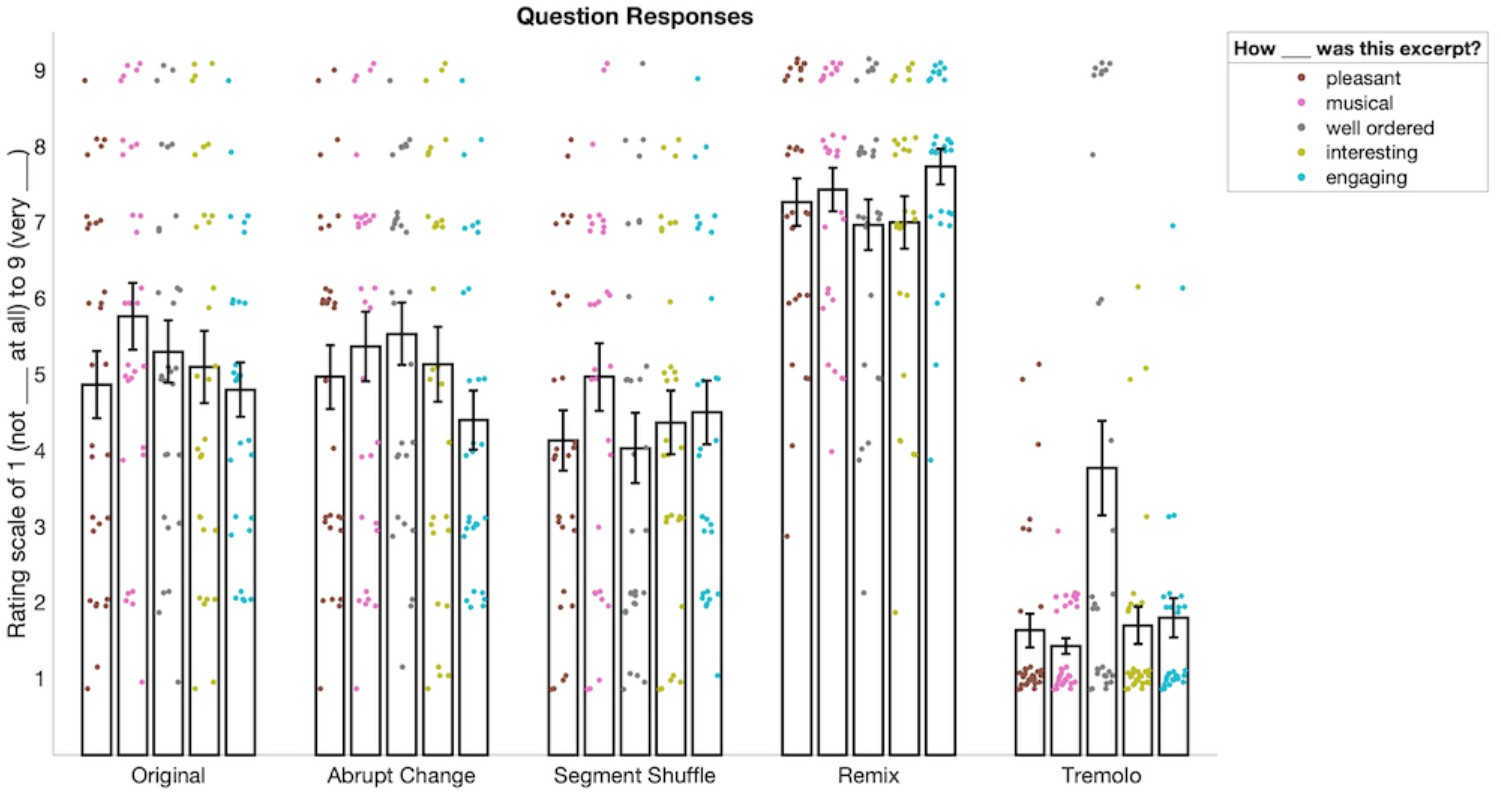
Behavioral ratings for all questions in the experiment (responses were ordinal and are slightly jittered for visualization only). Ratings for “pleasant”, “musical”, “well ordered”, and “interesting” come from Block 1 and ratings for “engaging” come from Block 2. For pleasant, musical, interesting, and engaging, responses for Remix were significantly higher than for the other conditions. For these same questions, responses were also significantly lower for Tremolo compared to all other conditions. For ratings of well ordered, we saw a similar pattern except that Abrupt Change was significantly higher than Segment Shuffle.

Follow-up pairwise t-tests comparing responses between conditions showed a similar pattern for four of the five questions (see Tables S2–S6 for all p-values). For pleasant, musical, interesting, and engaging ratings, Remix was significantly higher than the other four conditions (*p_FDR_* < 0.01, 10 comparisons) and Tremolo was significantly lower than the other four conditions (*p_FDR_* < 0.01). However, these ratings did not differ significantly between Original, Abrupt Change, and Segment Shuffle conditions.

Ratings for how “well ordered” the stimuli were followed a slightly different pattern. While Remix was significantly higher than all other conditions (see Table S4), Tremolo was significantly lower than all other conditions except Segment Shuffle (*p_FDR_* = 0.719). In addition, Segment Shuffle was significantly lower than Abrupt Change (*p_FDR_* = 0.036).

### 3.2 Full-Stimulus EEG ISC is Highest for Remix, Lowest for Tremolo

In computing the EEG ISCs, we first spatially filtered the responses for each stimulus in order to reduce their dimensionality from 125 electrodes to a single, maximally correlated spatial component (RC1) for each stimulus. These components are shown in Figure 5A. While our spatial filtering technique returned multiple components, we focus only on the first component because it is the only component with statistically significant coefficients for the majority of stimuli: Figure 5B demonstrates that RC1 was the only significant component for most stimuli (permutation testing; Original, Abrupt Change, Segment Shuffle, Remix *p_FDR_* < 0.001; Tremolo *p_FDR_* = 0.379; see Table S7 for all p-values).Remix also had a significant RC4 and Tremolo had no significant RCs. The topographies and coefficient significance for RC1 are in line with those computed in previous music EEG ISC studies (B. Kaneshiro et al., 2020, 2021); given that subsequent RCs did not correspond to significant ISC in a closely related study with similar distributions of coefficients (B. Kaneshiro et al., 2021), here we compute ISC only for RC1.

**Figure 5:**
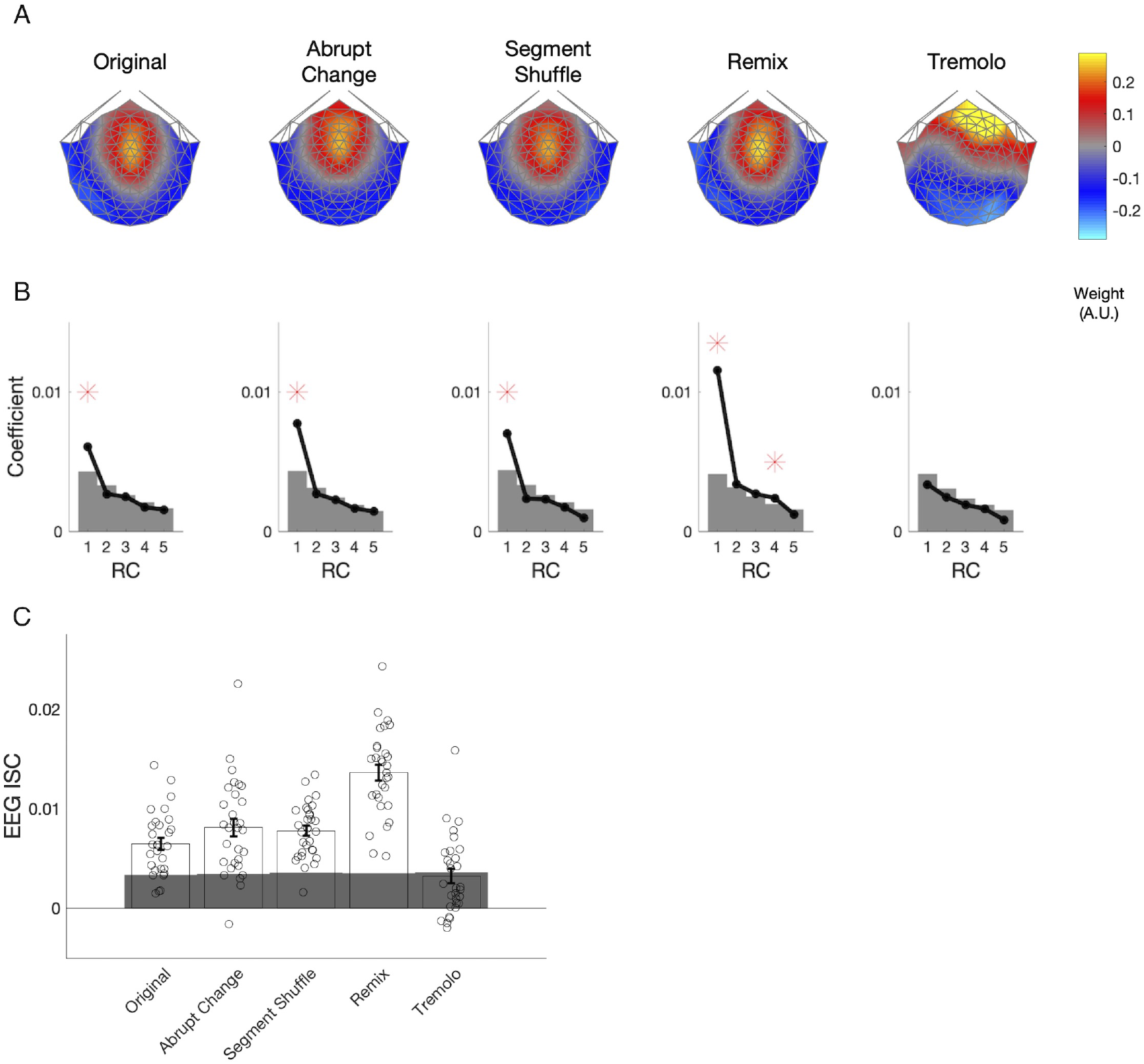
EEG components, coefficients, and aggregate ISC. (A) Spatial filter weights are visualized on a scalp model using forward-model projections. Maximally reliable components (RC1) exhibit consistent auditory topographies for all stimulus conditions except Tremolo. (B) Spatial filter eigenvalues serve as component coefficients. Significant coefficients are marked with red asterisks and significance thresholds; gray areas denote the 95th percentile of the null distribution. RC1 is statistically significant for all conditions except Tremolo. (C) ISC was computed over the entire duration of each stimulus. Remix elicited significantly higher ISC than all the other conditions, and Tremolo elicited significantly lower ISC than all other conditions.

When computed over the entire duration of a stimulus, EEG ISC differed significantly by condition (repeated-measures ANOVA, χ^2^(4) = 96.002, *p* < 0.001). Follow-up pairwise comparisons indicated that Original, Abrupt Change, Segment Shuffle, and Tremolo all significantly differed from Remix (*p_FDR_* < 0.001), and Original, Abrupt Change, Segment Shuffle, and Remix all differed from Tremolo (*p_FDR_* < 0.001). Figure 5C shows the direction of these significant differences: Remix garnered higher overall EEG ISC values than the other conditions, while Tremolo received the lowest overall values. Despite their structural differences, ISC among Original, Abrupt Change, and Segment Shuffle did not differ significantly from one another when computed over entire excerpts (see Table S8 for a full list of p-values).

### 3.3 Full-Stimulus CB ISC Alligns Broadly with EEG ISC

To analyze the CB ISC values (Figure 6), we followed the same procedures used for comparing EEG ISC computed over entire stimuli. These values significantly differed by condition (χ^2^(4) = 180.2, *p* < 0.001). Pairwise comparisons revealed that Remix had higher ISC than all other conditions, Tremolo had lower ISC than all other conditions, and Segment Shuffle had higher ISC than all conditions except Remix. All condition comparisons were significant except for Original versus Abrupt Change (*p_FDR_* = 0.87; all other comparisons, *p_FDR_* < 0.05, see Table S9 for a full list).

**Figure 6:**
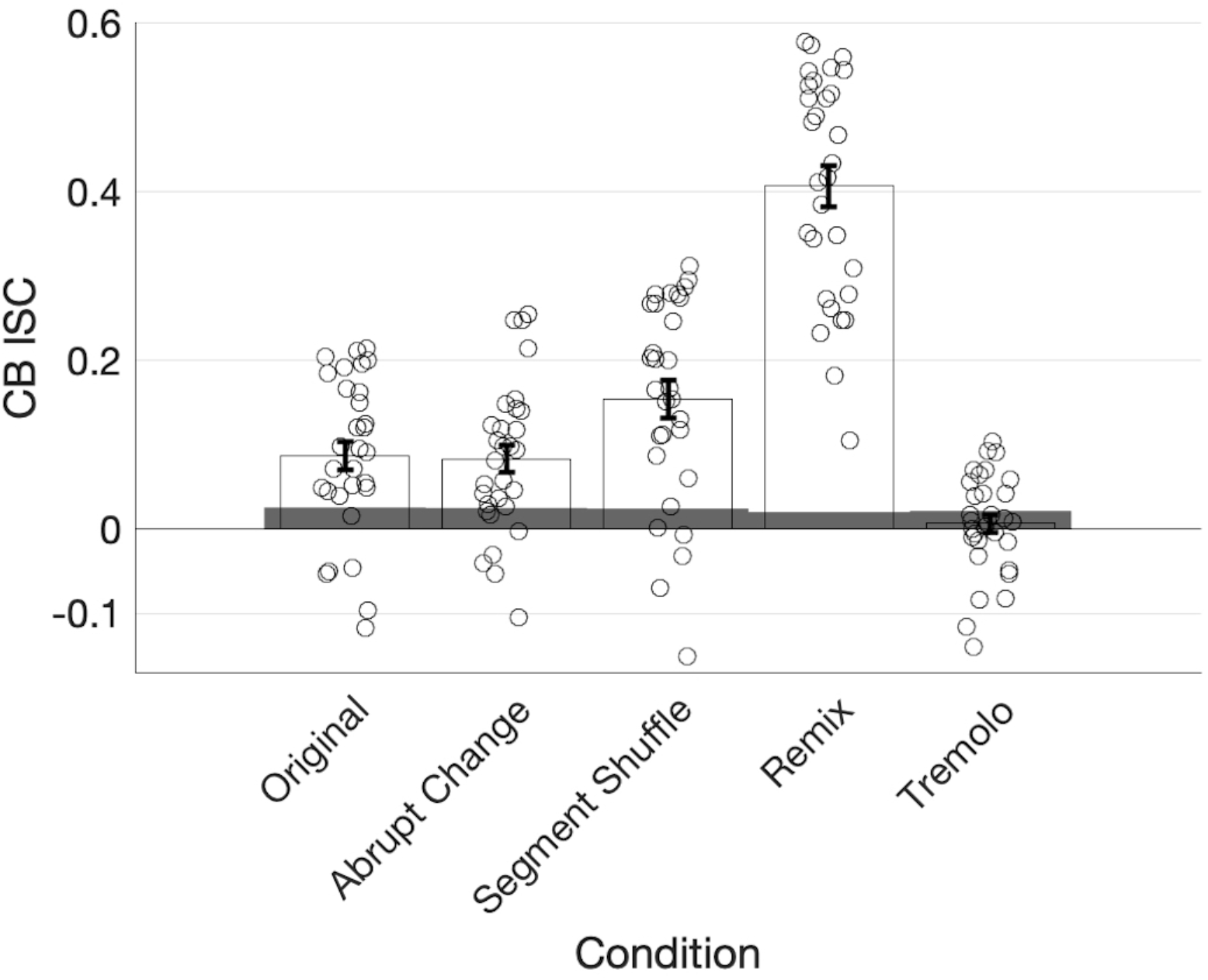
ISC of continuous behavioral (CB) reports of engagement for each condition with individual participant data and standard error of the mean plotted. Remix elicited significantly higher ISC than all the other conditions and Tremolo elicited significantly lower ISC than all the other conditions. Segment Shuffle also differs significantly from all other conditions.

### 3.4 Time-Resolved Measures Coincide with a Subset of Musical Events

In addition to calculating the overall ISC for EEG and CB data, we were interested in observing changes in ISC over the course of the stimuli. After computing ISC over short, shifting time windows, we visualized the ISC trajectory over time. Permutation testing provided a time-varying statistical significance threshold, allowing us to see when participants, as a group, delivered significantly correlated responses. Below we give a qualitative assessment of these results (Figure 7). Note that although EEG and CB ISC data had different sampling rates, we used identical time window lengths (5 seconds) and shifts (1 second) to facilitate comparison. We plot time-resolved ISC at the center of each temporal window. This means significant ISC implicates activity from ±2.5 seconds around each time point.

**Figure 7:**
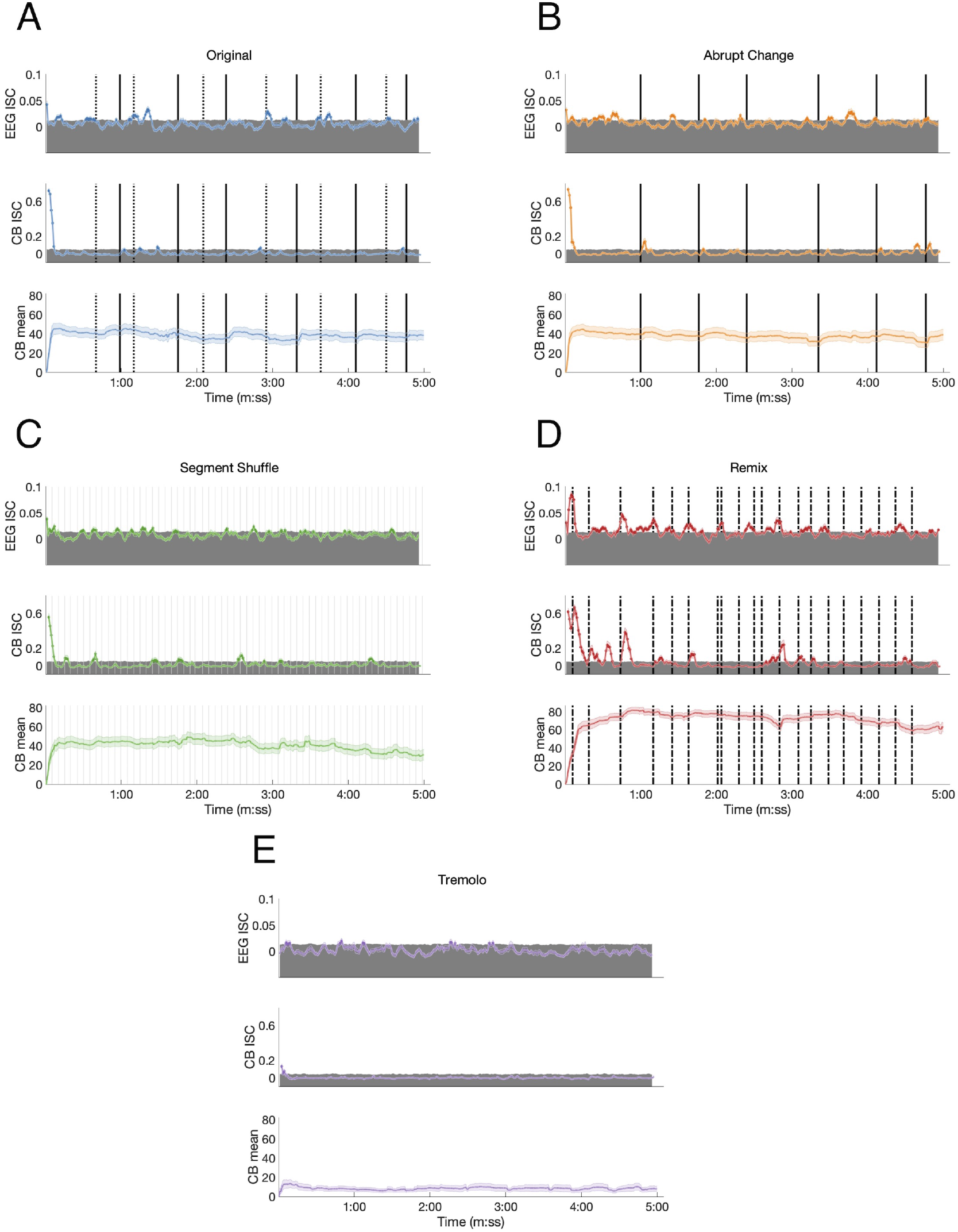
Time-resolved EEG ISC, CB ISC, and CB means for each condition. (A) Original: Dotted lines mark the start of phasing sections, solid lines mark the start of in-phase sections. (B) Abrupt Change: Solid lines mark the start of each new in-phase section. (C) Segment Shuffle: Light gray lines mark the start of each new segment. (D) Remix: Dashed lines mark musical events expected to be significant to listeners. (E) Tremolo.

Responses to the Original stimulus show small but significant ISC peaks in the EEG data (permutation test *p* < 0.05, uncorrected; see Methods), with statistically significant ISC in 16.9% of the time windows (Table 1). The largest ISC peaks appear around the approximate start times of phasing sections, or shortly thereafter. Each of the phasing section onsets (marked in Figure 7A with dotted lines) is accompanied by a significant peak with the exception of the third phasing section. While phasing elicits ISC peaks relatively consistently, in-phase sections fail to correspond to any significant ISC peaks. Both EEG and CB ISC also contain a significant peak at the start of the excerpt. In the time-resolved CB ISC data, only a handful of small peaks occur above the significance threshold after the initial drop; they seem unrelated to phasing and in-phase musical events, and only 4.7% of the ISC values are significant (Table 1). In contrast with phasing sections eliciting consistent peaks in the EEG ISC data, the CB mean data shows an increase in mean engagement rating after the start of each in-phase section. There also appears to be a slight decrease across the length of the stimulus.

**Table 1:** Percentages of significant time-resolved ISC for each condition for both EEG and CB data.

| Stimulus | % significant time-resolved EEG ISC | % significant time-resolved CB ISC |
| --- | --- | --- |
| Original | 16.892% | 4.65% |
| Abrupt Change | 18.581% | 6.98% |
| Segment Shuffle | 15.878% | 10.30% |
| Remix | 45.608% | 25.9% |
| Tremolo | 7.432% | 1.00% |

EEG ISC data for the Abrupt Change condition shows significant peaks within five seconds of the in-phase shifts (shifts number two, three, five and six as marked in solid lines in Figure 7B; (18.6% of ISC values are significant; see Table 1)). In contrast with the Original condition, in the Abrupt Change condition, where in-phase sections begin suddenly, they seem to elicit ISC peaks in the EEG data. The other small significance peaks in the EEG data come between in-phase changes, perhaps as participants anticipate stimulus alterations during the long stretches of unchanging material (perhaps something like the contingent negative variation between warning and imperative stimuli, (Tecce, 1972)). After an initial descent, the CB ISC data shows significant peaks around the first two and final two in-phase changes (percentage of significant time-resolved CB ISCs = 7.0%; see Table 1). The other two significant peaks appear between in-phase changes, perhaps related to the effect noted above. As in the Original condition, time-resolved CB mean data shows slight increases in engagement ratings after all six abrupt changes and an overall decline in engagement.

The perennially unpredictable changes in Segment Shuffle were met with frequent, small bursts of significant ISC correlations in the EEG data (Figure 7C; 15.9% significant ISC values; see Table 1). Comparing EEG and CB ISC time courses reveals unreliable alignment : After the initial drop in CB data, eight significant peak bursts unfold; about half of them align with EEG peaks (see peaks around time 1:30 and 3:05) while the other half do not (see peaks around time 0:15 and 2:30). CB means show small bumps in engagement ratings in the midst of a long-term downward trend (percentage of significant time-resolved CB ISCs = 10.3%; see Table 1).

Time-resolved ISCs for the Remix condition give ample opportunity to correlate peaks with musical events, with statistically significant EEG ISC in 45.6% time windows and significant CB ISC in 25.9% of time windows (Table 1). We selected the coded events in Figure 7D based on moments in the work that we deemed most musically salient (see Table S1 for the timings and descriptions of all twenty events). Note that not all of these events aligned with ISC peaks, but here we discuss some that did. After a sample from *Piano Phase* is presented for the first few seconds of Remix, a dramatic drum machine attack builds into simultaneous entrances for a synth countermelody and marimba riff (0:06). This build up and entrance align with the first and largest peak in the EEG data. The second peak in the EEG data comes at what might be the most dramatic moment in the piece, a beat drop anticipated with a drum machine lick (0:44). Note the potentially related peak in the CB ISC data following this event. But ISC peaks are not always elicited in both EEG and CB data. For example, the neighboring musical moments around minute 2:00 arise from a sudden dropping out of the percussion for a few seconds (2:01), leaving only a low, meandering synth line and a *Piano Phase* sample until the percussion reenters (2:04). This double event seems associated with an EEG ISC peak but no significant CB activity. A similar compositional technique plays out before minute 3:00. Two coded lines before that time (2:36), all instruments drop out except for the *Piano Phase* sample. It goes on, unchanging, until lush pitched percussion (a marimba) and additional synth lines enter at 2:50 (the line just before minute 3:00 in Figure 7D). The ISC peaks in both the EEG and CB data anticipate the reentry of additional instrumental lines, possibly in line with the previously mentioned effect: an anticipation that something must be coming given the static situation.

We did not expect any significant EEG ISC peaks for Tremolo, with its static, stark content. We see only occasional, small peaks above significance (Figure 7E; percentage of significant time-resolved EEG ISCs = 7.4%; percentage of significant time-resolved CB ISCs = 1.0%; see Table 1). We also note that in contrast to the other stimulus conditions, the time-resolved EEG ISC for this condition does not include a significant peak at the beginning of the excerpt. However, similar to the control condition in B. Kaneshiro et al. (2020), this RC1 differs in topography from the other conditions (Figure 5A) and is not statistically significant (Figure 5B), complicating interpretation of the ISC time course.

Comparing the present percentages of significant time-resolved ISCs for EEG data in RC1 with those reported by B. Kaneshiro et al. (2021) shows that our highest EEG ISC (for Remix) eclipses their finding of 37% (in response to Elgar’s cello concerto); our Original, Abrupt Change, and Segment Shuffle stimuli elicit higher percentages of significant ISC than their control condition (an envelope-scaled but otherwise temporally unstructured manipulation); and our Tremolo condition approximates the percentage found for their control condition (8%).

## 4 Discussion

We tested the limits of inter-subject correlation (ISC) as a measure of engagement with musical stimuli using Steve Reich’s *Piano Phase* as well as manipulated and remixed versions of the work. By comparing ISC results for EEG and continuous behavioral (CB) responses as well as behavioral ratings, we found no clear differences between manipulations based on compositional techniques (Original, Abrupt Change, and Segment Shuffle), but consistently high correlations and ratings for a popular-music version (Remix), and low correlations and ratings for a version featuring extreme repetition (Tremolo). These findings may underscore the subtlety of a core minimalist technique (phasing) and may also clarifies some limits of ISC as a measure of engagement with auditory stimuli.

The varied measures we collected (EEG, CB, behavioral ratings) align when viewing aggregate measures over entire stimuli. Aggregate EEG ISC and CB ISC, as well as behavioral ratings show Remix garnering significantly higher values than the other conditions, and Tremolo significantly lower values than the other conditions (see also the strong correlations between overall CB means and overall behavioral ratings of engagement in Figure S2). From this overall stance, EEG ISC values for Original, Abrupt Change, and Segment Shuffle do not differ from each other, and neither do participants’ behavioral ratings (with the single exception of ratings for “well ordered” mentioned above). Note that overall CB ISC has a slightly different pattern than EEG ISC, with CB ISC for Segment Shuffle pulling statistically ahead of Original and Abrupt Change. At the time-resolved level we notice differences between EEG ISC and other measures. Phasing sections in the Original, with their many and unpredictable onsets, elicit neural ISC but fail to generate CB ISC. Participants seemed to drift towards higher engagement ratings at the start of in-phase sections (see CB means), perhaps returning attention to the stimulus when it emerges from complex phasing sections back towards unison clarity (in-phase sections). We also noted the mix of alignment and independence between neural and behavioral measures in Remix, again with low-level acoustic changes attracting neural attention that elicits no behavioral ISC. Differences between EEG and CB ISC were also noted in Abrupt Change and Segment Shuffle conditions.

Previous studies have reported decreased ISC when music stimuli are repeated (Madsen et al., 2019; B. Kaneshiro et al., 2020). One explanation of our findings is that highly repetitive music (such as minimalism and Reich’s phasing process) will elicit lower engagement, and thus, lower ISC values. Certainly, our Tremolo condition offers an extreme test and seeming confirmation of this hypothesis. More varied stimuli still featuring high levels of repetition—i.e., Original, Abrupt Change, and Segment Shuffle—yielded higher EEG and CB ISC than Tremolo. Remix’s frequently changing musical parameters resulted in rather high ISC. One could argue that the more repetitive the stimulus was, the less interesting it may have been, and thus, less engaging.

Yet, as some have pointed out (Madsen et al., 2019; B. Kaneshiro et al., 2021), ISC measures *shared* engagement. Put another way, ISC can only pick up on forms of engagement that unfold similarly between multiple participants. Other types of engagement, be they idiosyncratic, or only shared by a few participants, would not show up. The strongest empirical evidence for such a view of our current data comes from individual CB responses (Figure S1). In said data, at least two participants (the highest two lines of raw data) show patterns of high and dynamic engagement in the Tremolo condition, a condition where we predicted and found very low EEG and CB ISC. Previous theoretical and empirical work bolsters the idea of multiple styles of engagement. The transportation and cognitive elaboration framework for engagement posit two strands of engagement: transportation, where audience members are locked into the content of the art object, tracking details; and cognitive elaboration, where an observer or listener is prompted by the stimulus to reflect on the artwork, drawing connections with other experiences and other knowledge (Green & Brock, 2000). David Huron’s listening styles offer even more potential types or modes of engagement, ranging from mentally singing along to mentally reminiscing about musically associated memories (Huron, 2002). ISC would be unlikely to pick up on these listening styles equally, and it would be odd if a single measure could.

Some cognitive science of music scholars have argued that repetition could augment individualized, internally focused experiences by gradually demanding less processing power and attention over time. Such a process may open up reflective space for listeners (Margulis, 2014). (This is in contrast with the type of engagement that might occur during dramatic moments like the beat drop in the first minute of Remix.) In *Piano Phase*, such a trajectory could be cyclical, with listeners drifting off into individual experience and tugged back into the details of the ongoing external stimulus events by changes in the music. If enough participants were drawn back to the stimulus details at the same time, neural responses could become sufficiently correlated to produce an ISC peak (perhaps something like the peak around minute 3:00 in the Original EEG ISC time-resolved data). In this line of thought, musicologists and music theorists have noted the long trajectories of expectation formation in minimalist music such as Reich’s. Cadences in tonal music often drive and ultimately resolve such expectations (what key are we in? where are we in the phrase? what harmonic and melodic activity is likely to come next?). Cadences and their accompanying harmonic trajectories are also present in minimalism but often in a stretched out form (Fink, 1996). Some listeners may lose interest along the way, while others may be drawn into granular detail and vary in what layer of granularity they are caught up in. Perhaps most move from state to state: For examples of the former situation, two participants in the present study noted that the Tremolo stimulus was difficult to listen to—“intense” in the words of one. Another participant stated that to them the stimuli were “all the same but with different layers.”

While our primary interest was engagement patterns in *Piano Phase*, this study was also motivated by a desire to clarify and delimit what EEG ISC may index. Previous literature has emphasized ISC as a measure of engagement, defined as “emotionally laden attention” (Dmochowski et al., 2012). A number of earlier findings raise questions about this relationship. Frequently and unexpectedly changing stimuli seem capable of driving correlated neural responses, perhaps pointing to a relationship between ISC and something like the orienting response (voluntary and automatic neural and behavioral responses to novel information, (E. Sokolov, 1990; E. N. Sokolov et al., 2002)). Dmochowski and colleagues reported relationships between EEG ISC and population ratings of Super Bowl commercials and found that an audio-visual stimulus with “repeated and jarring scene cuts” associated with “relatively strong neural reliability” drove ISC measures above population ratings (this stimulus was ultimately excluded in order to maintain stronger predictive performance of population ratings; (Dmochowski et al., 2014, Supplementary Note 3)). Ki et al. (2016) found that narratives in a foreign language elicited higher ISC than a narrative in the participants’ native language. Using two films as stimuli, Poulsen et al. (2017) reported a significant correlation between ISC and average luminance difference, suggesting that ISC for their primary component of interest “may indeed be driven by low-level visual evoked responses” (p. 5). Finally, B. Kaneshiro et al. (2020) noted that a stimulus manipulation in which measures of music were randomly re-ordered (and thus musically less meaningful but more surprising) resulted in higher EEG ISC than intact music. In the current experiment, the extreme musical parameters of minimalism, stimulus manipulations, and continuous behavioral ratings allowed us to further explore what ISC might index. If ISCs mark more cognitive-level, emotional engagement, manipulated stimuli with ostensibly less musical interest should result in lower ISC than purportedly more musically meaningful stimuli. This was not the case when comparing Original with Abrupt Change and Segment Shuffle conditions (though these three conditions each had significantly higher overall EEG and CB ISC compared with the Tremolo stimulus—the most extreme control). Additionally, we can compare neural ISC with continuous behavioral responses and overall ratings: alignment between measures could support the “engagement” interpretation of neural ISC. The overall ISC and behav-ioral ratings mostly reinforce three groups: Remix on top, Original, Abrupt Change, and Segment Shuffle in the middle (statistically undifferentiated), and Tremolo at the bottom. When examining time-resolved data, we saw a mixture of alignment and difference between EEG and CB ISC. Given the partial overlap, perhaps it is safest to say that, if we choose to use the term “engagement”, it may need qualification: Perhaps the type, kind, or style of engagement indexed by EEG ISC is more sensory biased and less cognitively driven than the word engagement usually connotes.

Given the scope of data used in ISC analyses and the complexity of the culturally embedded stimuli with which participants are interacting, testing limit cases such as minimalism helps draw bounds around the interpretation and appropriate deployment of ISC as a measure of engagement. It also reveals new layers of detail for scholars who work on the repertoire—a testing ground for theories of how the music can function for individuals. On that front, this study suggests important follow up research. For instance, alpha activity is thought to reflect meditative states (Lee et al., 2018). Therefore, alternative approaches to analyzing the EEG data—e.g., by assessing alpha power, or correlation thereof—may prove more appropriate measures for indexing listener states while listening to minimalist music. We might hypothesize that when participants are diversely engaged with a stimulus, a similar psychological state may be shared—but one that is better indexed by other means than EEG ISC. As alpha activity has been shown to index multiple states in varying locations, future research could also include interviews with music listeners to provide complementary insights into inter-individual differences in music listening. Such mixed-methods work could reveal patterns for calm versus bored listeners or time periods of boredom, interest, and relaxation.

## Author Contributions

TD, DN, JB, and BK designed the experiment. TD, NG, JB, and BK created the stimuli. DN and BK created participant interfaces for the experiment. TD and DN collected the data. TD, DN, and BK curated the data. TD, JD, and BK specified formal and statistical analyses. TD and BK analyzed the data. TD and BK created the visualizations. TD and BK drafted the original manuscript. DN, NG, JD, and JB reviewed and edited the manuscript. JB and BK supervised the research.

## Supplemental Data

The data generated and analyzed in this study can be found in the Naturalistic Music EEG Dataset—Minimalism (NMED-M) in the Stanford Digital Repository (https://purl.stanford.edu/kt396gb0630).

## Data Availability Statement

The data generated and analyzed in this study can be found in the Naturalistic Music EEG Dataset—Minimalism (NMED-M) in the Stanford Digital Repository (Dauer et al., 2021).^6^

## Supplementary Material

### 1 SUPPLEMENTARY DATA

The data generated and analyzed in this study can be found in the Naturalistic Music EEG Dataset—Minimalism (NMED-M) in the Stanford Digital Repository (https://purl.stanford.edu/kt396gbg630).

### 2 SUPPLEMENTARY FIGURES AND TABLES

#### 2.1 Figures

**Figure S1.**
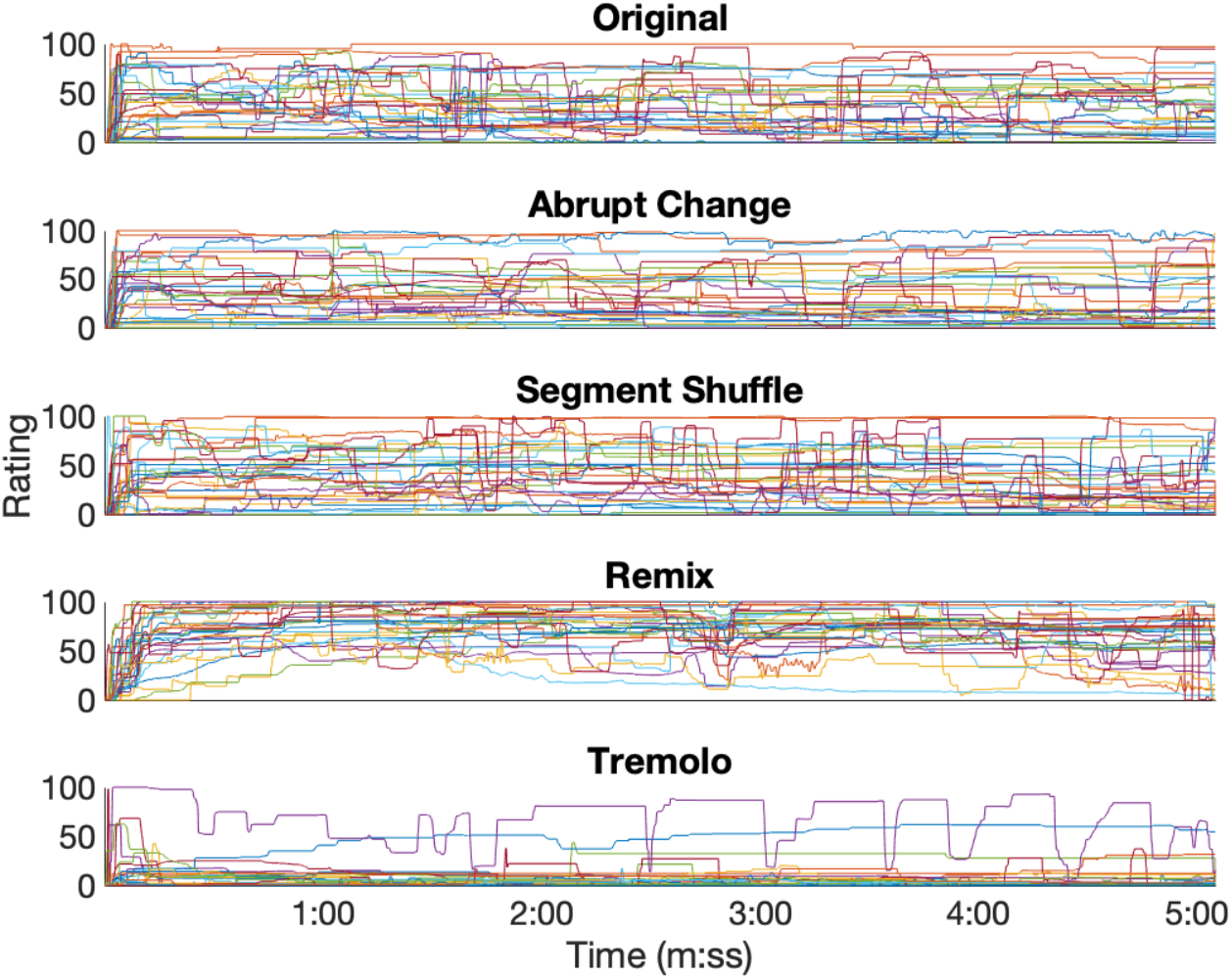
Continuous behavioral (CB) reports of engagement from individual participants, grouped by stimulus condition.

**Figure S2.**
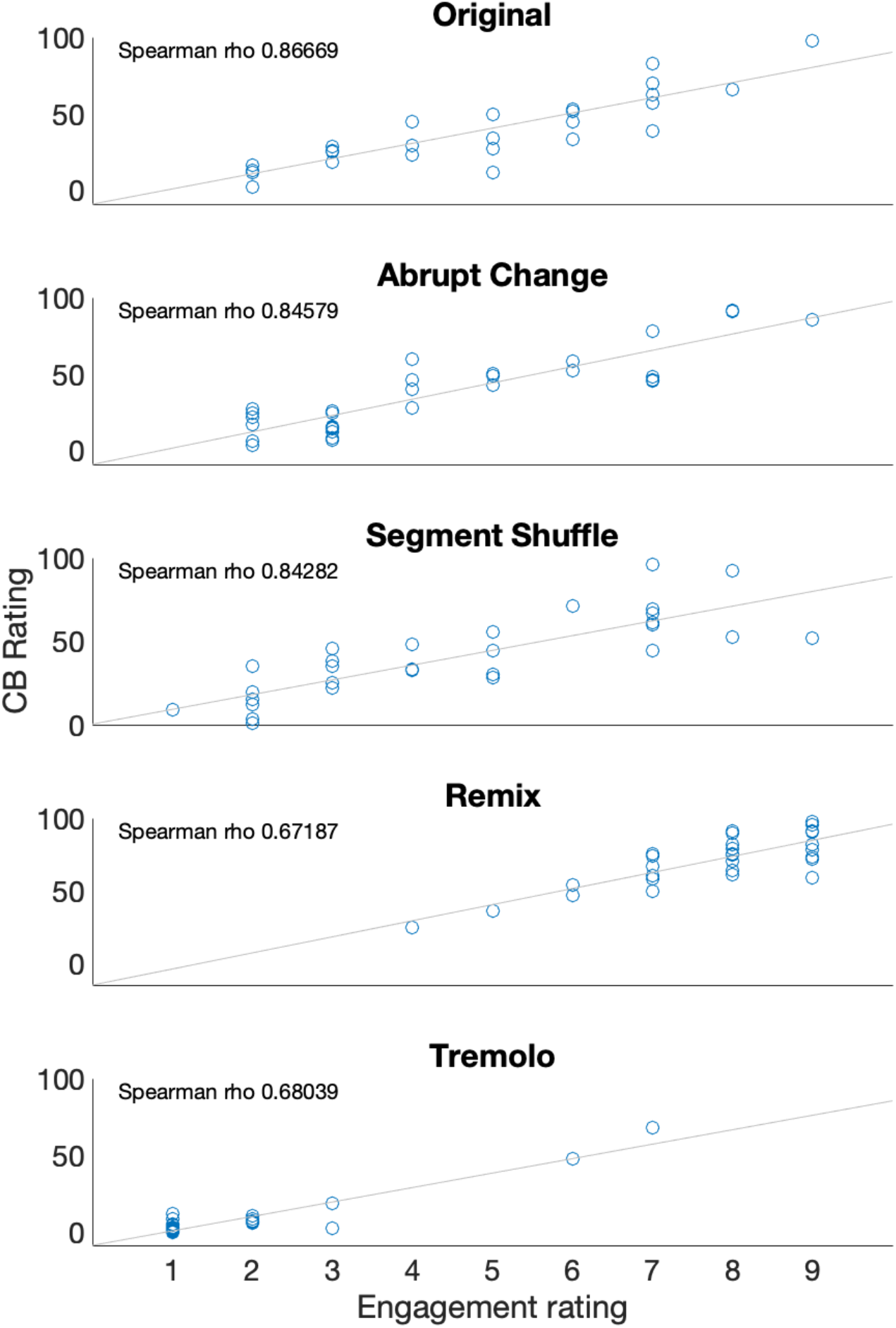
Scatter plot of each participant’s mean CB value with the behavioral rating in response to the question “How engaging was the stimulus?” Spearman’s rho also reported for each correlation.

#### 2.2 Tables

**Table S1.**
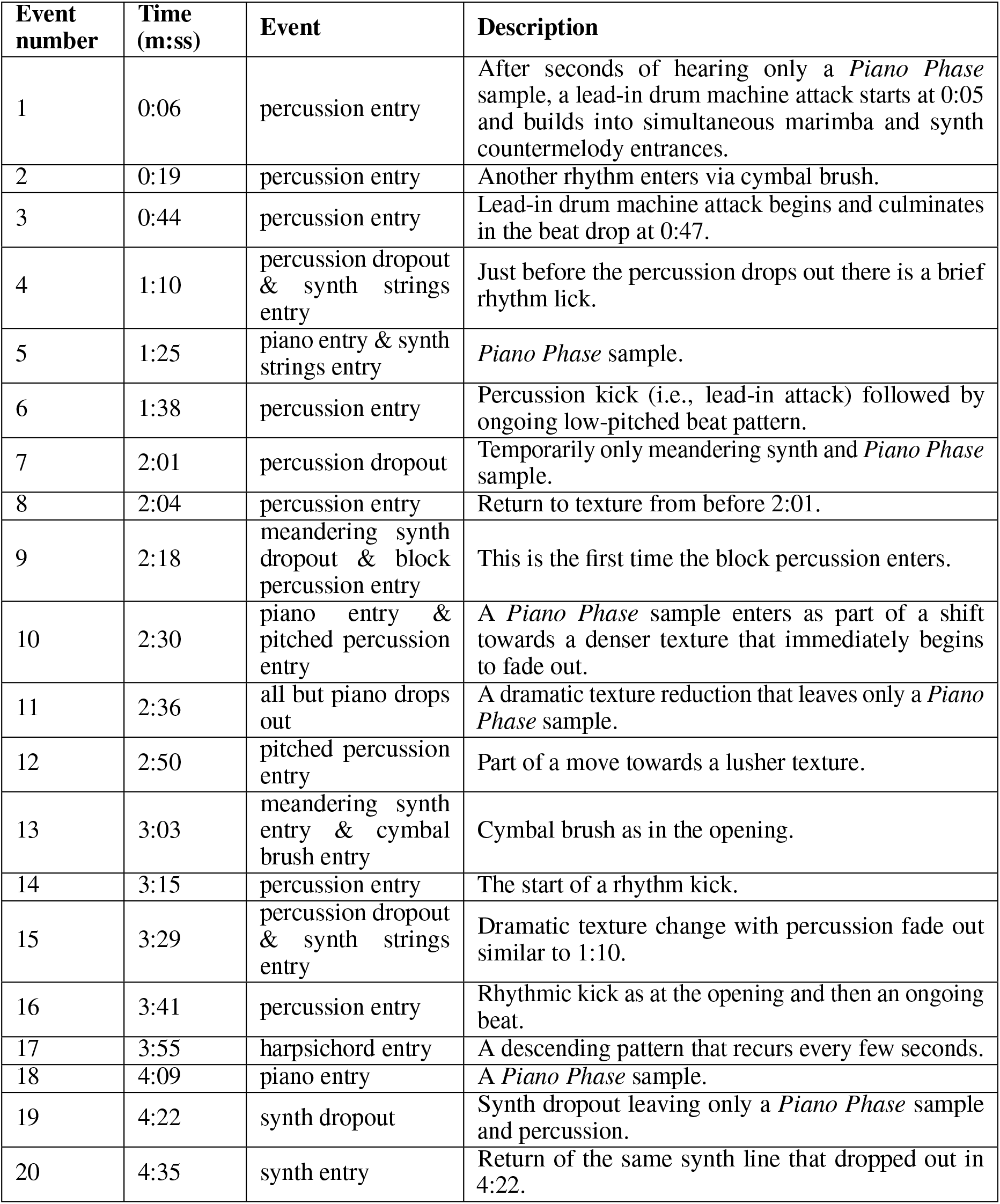
Musical events in Remix predicted to be salient for listeners.

**Table S2.** Post hoc t-test p-values (FDR-corrected) for ratings of how “pleasant” the stimulus was. Asterisks denote FDR-corrected p-values less than 0.05.

|  |  |  |  |  |
| --- | --- | --- | --- | --- |
| **Abrupt Change** | 0.848 |  |  |  |
| **Segment Shuffle** | 0.179 | 0.139 |  |  |
| **Remix** | < 0.001* | < 0.001* | < 0.001* |  |
| **Tremolo** | < 0.001* | < 0.001* | < 0.001* | < 0.001* |
|  | **Original** | **Abrupt Change** | **Segment Shuffle** | **Remix** |

**Table S3.** Post hoc t-test p-values (FDR-corrected) for ratings of how “musical” the stimulus was. Asterisks denote FDR-corrected p-values less than 0.05.

|  |  |  |  |  |
| --- | --- | --- | --- | --- |
| **Abrupt Change** | 0.449 |  |  |  |
| **Segment Shuffle** | 0.449 | 0.139 |  |  |
| **Remix** | 0.003* | < 0.001* | < 0.001* |  |
| **Tremolo** | < 0.001* | < 0.001* | < 0.001* | < 0.001* |
|  | **Original** | **Abrupt Change** | **Segment Shuffle** | **Remix** |

**Table S4.** Post hoc t-test p-values (FDR-corrected) for ratings of how “well ordered” the stimulus was. Asterisks denote FDR-corrected p-values less than 0.05.

|  |  |  |  |  |
| --- | --- | --- | --- | --- |
| **Abrupt Change** | 0.719 |  |  |  |
| **Segment Shuffle** | 0.065 | 0.036* |  |  |
| **Remix** | 0.027* | 0.040* | < 0.001* |  |
| **Tremolo** | 0.036* | 0.024* | 0.719 | < 0.001* |
|  | **Original** | **Abrupt Change** | **Segment Shuffle** | **Remix** |

**Table S5.** Post hoc t-test p-values (FDR-corrected) for ratings of how “interesting” the stimulus was. Asterisks denote FDR-corrected p-values less than 0.05.

|  |  |  |  |  |
| --- | --- | --- | --- | --- |
| **Abrupt Change** | 0.954 |  |  |  |
| **Segment Shuffle** | 0.226 | 0.226 |  |  |
| **Remix** | 0.002* | 0.002* | < 0.001* |  |
| **Tremolo** | < 0.001* | < 0.001* | < 0.001* | < 0.001* |
|  | **Original** | **Abrupt Change** | **Segment Shuffle** | **Remix** |

**Table S6.**
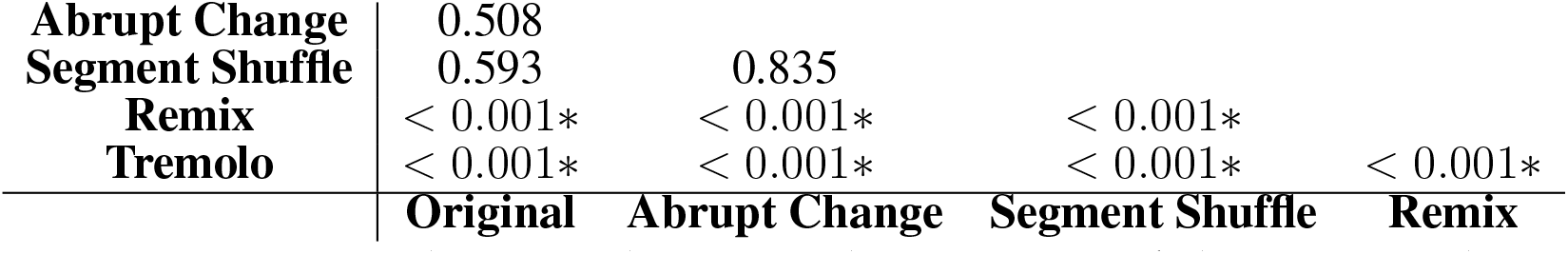
Post hoc t-test p-values (FDR-corrected) for ratings of how “engaging” the stimulus was. Asterisks denote FDR-corrected p-values less than 0.05.

**Table S7.** P-values (FDR-corrected) for RC coefficients. Asterisks denote FDR-corrected p-values less than 0.05.

|  | **Original** | **Abrupt Change** | **Segment Shuffle** | **Remix** | **Tremolo** |
| --- | --- | --- | --- | --- | --- |
| **RC1** | < 0.001* | < 0.001* | < 0.001* | < 0.001* | 0.379 |
| **RC2** | 0.442 | 0.379 | 0.723 | 0.079 | 0.442 |
| **RC3** | 0.231 | 0.259 | 0.379 | 0.050 | 0.442 |
| **RC4** | 0.433 | 0.379 | 0.379 | 0.005* | 0.379 |
| **RC5** | 0.238 | 0.194 | 0.761 | 0.442 | 0.836 |

**Table S8.**
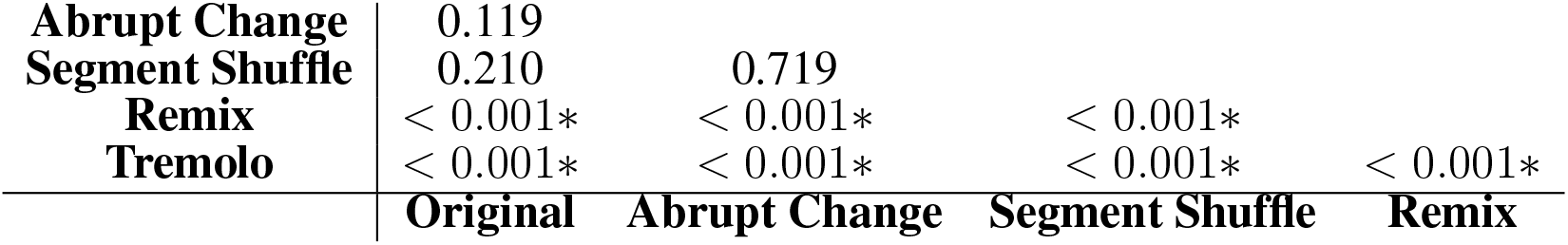
Post hoc t-test p-values (FDR-corrected) for overall EEG ISC values. Asterisks denote FDR-corrected p-values less than 0.05.

|  |  |  |  |  |
| --- | --- | --- | --- | --- |
| **Abrupt Change** | 0.119 |  |  |  |
| **Segment Shuffle** | 0.210 | 0.719 |  |  |
| **Remix** | < 0.001* | < 0.001* | < 0.001* |  |
| **Tremolo** | < 0.001* | < 0.001* | < 0.001* | < 0.001* |
|  | **Original** | **Abrupt Change** | **Segment Shuffle** | **Remix** |

**Table S9.** Post hoc t-test p-values (FDR-corrected) for overall CB ISC values. Asterisks denote FDR-corrected p-values less than 0.05.

|  |  |  |  |  |
| --- | --- | --- | --- | --- |
| **Abrupt Change** | 0.874 |  |  |  |
| **Segment Shuffle** | < 0.001* | < 0.001* |  |  |
| **Remix** | 0.010* | 0.014* | < 0.001* |  |
| **Tremolo** | 0.007* | 0.005* | < 0.001* | < 0.001* |
|  | **Original** | **Abrupt Change** | **Segment Shuffle** | **Remix** |

## Footnotes

1 The piece begins with one pianist (Pianist 1) playing a twelve-note pattern consisting entirely of sixteenth notes and containing five unique pitches in the treble register. The pattern can be divided into two groups of six sixteenth notes, and Reich gave a metronome marking of 72 beats per minute to the dotted quarter note (one group of six sixteenth notes). The score consists of numbered modules that are repeated an indeterminate number of times: Reich noted approximate ranges for the number of repetitions above each module. After the pattern is established in the first module, the second pianist (Pianist 2) fades in, playing the identical pattern in unison with Pianist 1. After repeating the pattern in unison for some time, Pianist 2 accelerates very slightly while Pianist 1 holds the opening tempo, causing the sound from the two pianos to wobble out of sync to varying degrees as the pattern is repeated at different tempos (we call these portions *phasing* sections). Various and unpredictable rhythm and pitch events emerge and disappear in these phasing sections. Eventually Pianist 2’s acceleration process culminates in another unison module where each pianist’s sixteenth notes are once again realigned (which we label *in-phase* sections). While the pianists’ rhythms are realigned, the pitch content of the pattern will have shifted: In this example, Pianist 2 aligns the second pitch of the opening pattern with the first pitch of the pattern (played by Pianist 1). *Piano Phase* proceeds by alternating between phasing and in-phase sections, where each successive in-phase section presents the next shifted alignment of the opening, twelve-note pattern (note three aligns with the first note of the pattern, a phasing section occurs, then note four aligns with the first note of the pattern, etc.).

2 We note that Madsen et al. (2019) did include Philip Glass’s *String Quartet No. 5* (1991): a more popular or “post-minimalist” work by comparison.

3 https://maplelab.net/reich/

4 A meter shift and accompanying pattern change occur later in the piece, but after the excerpt used in the experiment.

5 https://www.mattwinn.co.uk/about

6 Available at https://purl.stanford.edu/kt396gb0630.

